# MicroRNA-194 promotes lineage plasticity in advanced prostate cancer

**DOI:** 10.1101/752709

**Authors:** Rayzel C. Fernandes, John Toubia, Scott Townley, Adrienne R. Hanson, B. Kate Dredge, Katherine A Pillman, Andrew G. Bert, Richard Iggo, Rajdeep Das, Daisuke Obinata, MURAL investigators, Shahneen Sandhu, Gail P. Risbridger, Renea A. Taylor, Mitchell G. Lawrence, Lisa M. Butler, Amina Zoubeidi, Philip A. Gregory, Wayne D. Tilley, Theresa E. Hickey, Gregory J. Goodall, Luke A. Selth

**Affiliations:** Dame Roma Mitchell Cancer Research Laboratories, Adelaide Medical School, The University of Adelaide, Adelaide, South Australia, Australia; Freemasons Foundation Centre for Men’s Health, Adelaide Medical School, The University of Adelaide, Adelaide, South Australia, Australia; ACRF Cancer Genomics Facility, Centre for Cancer Biology, An alliance of SA Pathology and University of South Australia, Frome Road, Adelaide, South Australia, Australia; Centre for Cancer Biology, An alliance of SA Pathology and University of South Australia, Adelaide, South Australia, Australia; Institut Bergonié Unicancer, INSERM U1218, Bordeaux, France; Department of Radiation Oncology, University of California San Francisco, San Francisco, California; Department of Urology, Nihon University School of Medicine, Tokyo, Japan; Department of Anatomy and Developmental Biology, Monash Partners Comprehensive Cancer Consortium, Monash Biomedicine Discovery Institute, Prostate Cancer Research Group, Monash University, Clayton, Victoria, Australia; Cancer Research Program, Cancer Research Division, Peter MacCallum Cancer Centre, University of Melbourne, Melbourne, Victoria, Australia; Sir Peter MacCallum Department of Oncology, The University of Melbourne, Parkville, Victoria, Australia; Department of Physiology, Monash Partners Comprehensive Cancer Consortium, Monash Biomedicine Discovery Institute, Prostate Cancer Research Group, Monash University, Clayton, Victoria, Australia; South Australian Health and Medical Research Institute, Adelaide, South Australia, Australia; The Vancouver Prostate Centre, University of British Columbia, Vancouver, British Columbia, Canada; Faculty of Health and Medical Sciences, The University of Adelaide, Adelaide, South Australia, Australia; School of Biological Sciences, The University of Adelaide, Adelaide, South Australia, Australia

## Abstract

MicroRNA-194 (miR-194) promotes prostate cancer metastasis, but the precise molecular mechanisms by which it achieves this are unknown. Here, by integrating Argonaute high-throughput sequencing of RNA isolated by crosslinking immunoprecipitation (Ago-HITS-CLIP) with RNA sequencing and exon-intron split analysis, we defined a 163-gene miR-194 “targetome” in prostate cancer. These target genes were predominantly down-regulated through canonical 3’UTR recognition sites and were enriched within pathways involved in cytoskeletal organisation and cell movement. In clinical prostate cancer samples, miR-194 activity was inversely correlated with the androgen receptor (AR) signalling axis. At a mechanistic level, this inverse correlation was explained by down-regulation of miR-194 expression by AR. Accordingly, miR-194 expression and activity was significantly elevated in neuroendocrine prostate cancer (NEPC), an aggressive AR-independent disease subtype. MiR-194 enhanced the transdifferentiation of prostate adenocarcinoma cells to a neuroendocrine-like state, at least in part by targeting FOXA1, a transcription factor with a key role in maintaining the prostate epithelial lineage. Importantly, a miR-194 inhibitor effectively inhibited the growth of cell lines and patient-derived organoids with neuroendocrine features. Overall, our study reveals a novel post-transcriptional mechanism regulating the plasticity of prostate cancer cells and provides a rationale for targeting miR-194 in this NEPC.

## INTRODUCTION

Cellular plasticity, also referred to as lineage plasticity or lineage switching, is a process whereby cells exhibit reversible changes in properties and phenotypes. Cancer cells exploit this phenomenon in response to a targeted therapy, acquiring the phenotypic characteristics of another lineage that does not depend on the drug target for survival (1). This phenomenon allows cancer cells to adapt to new or stressful conditions and is increasingly recognised as a key feature of cancer progression (2).

As first-line treatment for metastatic prostate cancer (PCa), androgen deprivation therapy targets the exquisite dependence of tumours on the androgen receptor (AR) for their growth. While initially effective, patients inevitably develop resistance and progress to castration-resistant prostate cancer (CRPC). Most CRPC tumours exhibit adaptive changes that maintain AR activity despite the low androgen environment, an understanding that led to the development of highly potent second-generation AR-targeted therapies (e.g. Enzalutamide and Abiraterone). However, response to these newer agents is also limited in most cases (3). It has become increasingly clear that prolonged targeting of the AR, particularly with the more potent second-generation therapies, can drive cellular plasticity in CRPC. This plasticity is characterised by cells losing dependence on AR and gaining new phenotypes (i.e. aggressive variant PCa), with the most well recognised of these being a neuroendocrine (NE)-like state that is characterised by the expression of neuroendocrine, neuronal, developmental and stem cell markers (1). Neuroendocrine prostate cancer (NEPC) is evident in ∼15-25% of CRPC tumours (4,5) and exhibits aggressive clinical features; indeed, patients with NEPC have a median overall survival time of <1 year (4). A deeper understanding of how AR-targeted therapies promote lineage plasticity and the emergence of aggressive disease phenotypes such as NEPC is essential to improve patient outcomes.

Genomic comparisons of NEPC and CRPC adenocarcinoma (CRPC-Adeno) have revealed surprisingly few genetic differences between these disease subtypes; reproducible alterations in NEPC include higher incidences of *RB1* and *TP53* loss and more frequent amplification of *MYCN* and *AURKA* (6,7). The similarities in mutational landscapes between NEPC and CRPC-Adeno suggest that the plasticity underlying transdifferentiation from adenocarcinoma to an NE-like state is predominantly mediated by changes in epigenetics, transcriptional programs and protein function in the tumour cells, as opposed to selection and outgrowth of rare genetic variants (1).

In this study, we identified miR-194 as a novel post-transcriptional regulator of transdifferentiation in PCa. By targeting genes that suppress plasticity, such as *FOXA1*, miR-194 drives the emergence and growth of NEPC, a finding that justifies further investigation of miRNA-based therapies for this aggressive CRPC subtype.

## MATERIAL AND METHODS

### Cell lines and cell culture

LNCaP, PC3 and 22RV1 cell lines were purchased from the American Type Culture Collection (ATCC). LNCaP-MR42D and LNCaP-MR49F cell lines have been described previously (8). LNCaP and 22RV1 cell lines were maintained in RPMI-1640 (Sigma) containing 10% Fetal Bovine Serum (FBS) (Sigma). PC3 cell lines were maintained in RPMI-1640 containing 5% FBS. LNCaP-MR42D and LNCaP-MR49F cells were maintained in RPMI-1640 containing 10% FBS and 10μM Enzalutamide. For serum starvation experiments, cells were grown in phenol red-free RPMI-1640 containing 10% dextran-coated charcoal (DCC) stripped serum. Cell lines were subjected to regular mycoplasma testing. All cell lines underwent verification by short tandem repeat profiling by CellBank Australia.

### Cell line transfections

Transient transfection of cell lines were performed using RNAiMAX Transfection Reagent (Life Technologies) according to the manufacturer’s instructions. For HITS-CLIP and RNA-seq experiments, 22RV1 cells were transfected with 20nM miRVana mimic (miR-194 or negative control; Ambion). For all other experiments, cells were transfected with 20nM miRNA mimics from Shanghai GenePharma. For miR-194 inhibition, cells were transfected with 12.5 or 6.25nM locked nucleic acid (LNA) miRNA inhibitors (miR-194 LNA inhibitor or negative control inhibitor; Qiagen).

### Argonaute high-throughput sequencing of RNA isolated by crosslinking immunoprecipitation (Ago-HITS-CLIP)

The Ago-HITS-CLIP method was adapted from published methods (9,10), incorporating modifications from eCLIP (11,12). 22RV1 cells were seeded in 10 cm cell culture dishes and transfected in suspension with 20 nM miRVana mimic (miR-194 or negative control, 3 replicates of each; Ambion) using RNAiMAX (Life Technologies). After 24 h, transfected 22RV1 cells were rinsed once with ice-cold PBS and UV irradiated with 600 mJ/cm2, 254 nm, in ice-cold PBS using a UV Stratalinker-1800 (Agilent). Cells were collected by scraping, and cell pellets stored at -80°C as one pellet per 100mm plate. One pellet per CLIP IP was lysed in 500 μl of 1 X PXL (1 X PBS, 0.1% SDS, 0.5% deoxycholate, 0.5% Igepal) + EDTA-free Complete protease inhibitor cocktail (PIC; Roche) for 15 min on ice, followed by trituration through a 21G needle and syringe 5 times. DNA was digested with 20 μl RQ1 DNAse (Promega) at 37°C for 10 min on a Thermomixer (750 rpm, Eppendorf). RNA was partially digested with RNase 1 (ThermoFisher) by adding 5 μl of 1:40 diluted RNase 1 in 1 X PBS at 37°C for 5 min on a Thermomixer (750 rpm), then returned to ice. Lysates were centrifuged at 21,000 g for 20 min at 4°C and supernatant transferred to a fresh tube.

AGO-RNA complexes were immunoprecipitated using mouse IgA2 monoclonal anti-Ago2 antibody 4F9 (13); hybridoma sourced from University of Florida ICBR) with a mouse IgA antibody (GeneTex S107) used as a control. Antibodies (8 μg) were conjugated to 20 μl protein L Dynabeads (ThermoFisher, 88849) in PBS-Tw (1 X PBS, 0.05% Tween-20) for 45 min and washed three times with 1 X PXL (1 X PBS, 0.1% SDS, 0.5% sodium deoxycholate, 0.5% Igepal) before resuspending the beads with 450 μl of prepared lysate and rotating for 2 hr at 4°C. Bound AGO-RNA complexes were washed twice each consecutively with ice cold 1 X PXL, 5 X PXL (5 X PBS, 0.1% SDS, 0.5% sodium deoxycholate, 0.5% Igepal), and 1 X PNK (50 mM Tris-Cl pH 7.5, 10 mM MgCl2, and 0.5% Igepal). Beads were first treated with T4 PNK (NEB, M0201L; 20 U in 80 μl reaction volume) in the absence of ATP (37°C, 850 rpm for 20 min) to dephosphorylate 3’ RNA ends followed by washes with 1 X PNK, 5 X PXL, and two washes with 1 X PNK at 4°C. The 3’ preadenylated linker (NEBNext 3’SR adaptor for Illumina; /5rApp/AGA TCG GAA GAG CAC ACG TCT /3AmMO/) was ligated to the RNA fragments on bead using T4 RNA ligase 2 truncated KQ (NEB M0373; 100 U in a 40 μl reaction volume, 12% PEG8000, 1x RNA ligase buffer, 0.125 μM adaptor) in the absence of ATP at 16°C, overnight with periodic mixing. Beads were washed consecutively with ice cold 1 X PXL, 5 X PXL, and twice with 1 X PNK. Bound RNAs were then labelled with P32 γ-ATP using T4 PNK, 20 min at 37°C, and washed as above.

AGO-RNA complexes were eluted with 40 μl 1 X Bolt LDS sample buffer (ThermoFisher) + 1% β-mercaptoethanol at 70°C for 10 min on a Thermomixer (1200 rpm). Samples were separated through Bolt 8% Bis-tris Plus gels (ThermoFisher) using BOLT MOPS SDS running buffer at 200 V for 75 min. Complexes were then transferred to nitrocellulose (Schleicher&Schuell, BA-85) by wet transfer using 1 X BOLT transfer buffer with 10% methanol. Filters were placed on a phosphor screen and exposed using a Typhoon imager (GE). 115-160 kDa regions (corresponding to RNA tags > 30 nt) were excised from the nitrocellulose. RNA was extracted by proteinase K digestion (2 mg/mL proteinase K, 100 mM Tris-HCl pH 7.5, 50 mM NaCl, 10 mM EDTA, 0.2% SDS) at 50°C for 60 min on a Thermomixer (1200 rpm) followed by extraction with acid phenol (ThermoFisher, AM9712) and precipitation with 1:1 isopropanol:ethanol. RNA was pelleted by centrifugation then separated on a 15% denaturing polyacrylamide gel (1:19 acrylamide, 1 X TBE, 7 M urea). The wet gel was wrapped in plastic wrap and exposed to a phosphor screen and imaged using a Typhoon. Gel slices were cut corresponding to the expected size of the cross-linked RNA eluted by the “crush and soak” method as previously described (10).

Reverse transcription, 5’ linker ligation and amplification were performed essentially as previously described (11) using SR-RT primer for reverse transcription (IDT, AGACGTGTGCTCTTCCGATCT) with SuperScript IV, and a custom synthesised 5’ linker (IDT, 5’SRdeg /5Phos/NN NNN NNN NNG ATC GTC GGA CTG TAG AAC TCT GAA C/3SpC3/). Products were amplified for 20 cycles using a common forward primer (NEBNext SR primer for Illumina) and barcoded reverse primers for each sample (NEBNext Index primers for Illumina). PCR products were purified using Qiagen Qiaquick PCR purification kit, separated on an 8% acrylamide (29:1) TBE non-denaturing gel, stained with SYBR Gold nucleic acid gel stain (ThermoFisher) and imaged on a ChemiDoc (BioRad). Products corresponding to an insert size of ∼30 – 70 nt were excised from the gel and extracted by the “crush and soak” method as previously described (10). Library quality and quantity was assessed by Bioanalyzer (Agilent) and qPCR, pooled and sequenced on an Illumina NextSeq 500 (1 × 75bp).

RNA libraries generated by HITS-CLIP were sequenced on the Illumina Nextseq 500 platform using the single end protocol with a read length of 75. Raw reads were adapter trimmed and filtered for short sequences using cutadapt v1.8.1 (14) setting minimum-length option to 18, error-rate 0.2, and overlap 5. The resulting FASTQ files (averaging 41.6 million reads per sample) were analysed and quality checked using the FastQC (http://www.bioinformatics.babraham.ac.uk/projects/fastqc) program. Filtered reads were mapped against the human reference genome (hg19) using the Tophat2 alignment algorithm (version 2.0.9 with default parameters) (15), returning an average alignment rate of 43.8%. Unique molecular identifiers (UMIs) were used to de-duplicate reads that mapped to the same start site, possessed identical CIGAR strings and UMI barcodes sequences ≤1 edit distance apart. Enriched regions of the genome were identified from Samtools quality-filtered alignments (16) (-q 5) with the MACS2 peak caller (version 2.1.1) (17) (setting; --nomodel, --shift -15, --extsize 50, -B, --slocal 0, --llocal 0, --fe-cutoff 10, -q 0.05). Peak calling was performed using pooled alignment files and carried out separately for each strand. The resulting peak files from each strand were merged. Features in the vicinity of peak loci and enrichment of motifs within peaks were determined and analysed using Homer (18). Alignments were visualised and interrogated using IGV (19).

CLIP using a control antibody was performed on a single biological replicate of control transfected cells but yielded very little sequence data and was excluded from the analysis.

### RNA sequencing

22RV1 cells were seeded in 6-well plates and transfected in solution with 20nM miRVana mimic (miR-194 or negative control; Ambion) using RNAiMAX (Life Technoloies). At 36 hours post-transfection, cells were collected in Qiazol (Qiagen) and total RNA was extracted using a miRNeasy Mini Kit (Qiagen) according to the manufacturer’s instructions. RNA seq was performed on 4 biological replicates each of 22RV1 cells transfected with miR-194 or negative control. RNA sequencing libraries were constructed with the mRNAseq Library prep kit and libraries were sequenced on the Illumina NextSeq 500 platform.

RNA-seq libraries were multiplexed and sequenced on the Illumina NextSeq 500 platform using the stranded, paired-end protocol with a read length of 150. Raw reads were adapter trimmed and filtered for short sequences using cutadapt v1.8.1 (14), setting minimum-length option to 18, error-rate 0.2, quality cut-off 28, overlap 5 and trim N’s on. The resulting FASTQ files (averaging 60.2 million read pairs per sample) were analysed and quality checked using the FastQC program (http://www.bioinformatics.babraham.ac.uk/projects/fastqc). Reads were mapped against the human reference genome (hg19) using the STAR spliced alignment algorithm (20) (version 2.5.3a with default parameters and --chimSegmentMin 20, --quantMode GeneCounts), returning an average unique alignment rate of 92.9%. Differential expression analysis was evaluated from TMM normalised gene counts using R (version 3.2.3) and edgeR (21) (version 3.3), following protocols as described (22). Graphical representations of differentially expressed genes were generated using Glimma (23). Alignments were visualised and interrogated using the Integrative Genomics Viewer v2.3.80 (19).

Exon Intron Split analysis (EISA) was performed as described previously (24). To refine the miR-194 targetome, only post-transcriptionally downregulated genes (i.e. genes with log2FC(dExon-dIntron) < 0) and a FDR cutoff of 0.05) were considered as targets.

### Gene set enrichment analysis

Genes were ranked according to expression using the Signal2Noise metric. Gene Set Enrichment Analysis (Preranked analysis) (25) was implemented using the Broad Institute’s public GenePattern server with default parameters.

### Analysis of miR-194 activity in published datasets single sample GSEA (ssGSEA)

Expression data was downloaded from GEO (Kumar 2016 (GSE77930) (26)), cBioportal (MSKCC (27) and SU2C (28)), TCGA (29) and dbGAP (Beltran 2016 (30)). ssGSEA (31) was implemented using the Broad Institute’s public GenePattern server, using rank normalisation and default parameters. Since miRNAs repress expression of their target genes, miR-194 activity was calculated as the inverse value of ssGSEA scores for the miR-194 targetome.

### RNA extractions from cell lines and patient-derived xenograft (PDX) tissues

Total RNA from cell lines was extracted using TRI Reagent (Sigma), as described previously (32). PDX tissues preserved in RNALater were provided by the Melbourne Urology Research Alliance (MURAL) (33). Tissues were homogenised in Qiazol (Qiagen) with a Precellys24 Tissue Homogeniser (Bertin Technologies) and total RNA was extracted using a miRNeasy Mini Kit (Qiagen), according to the manufacturer’s instructions.

### Quantitative RT-PCR (qRT-PCR) analysis of mRNA

Total RNA was treated with Turbo DNA-free kit (Invitrogen), and reverse transcribed using iScript Reverse Transcriptase Supermix kit (Bio-Rad). qRT-PCR was performed in triplicate as described previously (34). GAPDH levels were used for normalization of qRT-PCR data. Primer sequences are available on request.

### qRT-PCR analysis of miR-194

Total RNA (100 ng) was reverse transcribed using the TaqMan MicroRNA Reverse Transcription Kit (Applied Biosystems) and Taqman Microarray Assays (Thermo Fisher Scientific). Quantitation of miR-194, U6 and RNU44 was done by qRT-PCR using Taqman Microarray Assays (Thermo Fisher Scientific) and TaqMan™ Universal Master Mix II, no UNG (Applied Biosystems) on a CFX384 real-time PCR detection system (Bio-Rad). MiR-194 expression was normalised to expression of U6 (cell lines) or the geometric mean of U6 and RNU44 (PDX tissues).

### Proliferation and cell viability assays

Proliferation curves for cell lines treated with LNA miRNA inhibitors were performed using the Trypan blue exclusion assay. Cells were seeded at 1×10^4^ (PC3) or 4.5×10^4^ (LNCaP-MR42D, LNCaP-MR49F, LNCaP) in 12-well plates and transfected in suspension with 12.5 or 6.25 nM miR LNA inhibitor using RNAiMAX (Life Technologies). Live and dead cells were quantified at indicated time points using Trypan blue.

For cell viability assays, LNCaP-MR42D or LNCaP cells were seeded at 4×10^3^ cells/well in 96-well plates and transfected in suspension with 12.5 or 6.25nM miR LNA inhibitor using RNAiMAX (Life Technologies). Cell viability was assesses using the Cell Titer-Glo Luminescent Cell Viability Assay (Promega) according to manufacturer’s recommendations.

### Neurite length measurement

Length of neurite extensions were measured using the Simple Neurite Tracer plugin (35) for Fiji/ImageJ. Neurite lengths were measured from ≥ 3 images per replicate. Representative images with overlaid traces were generated using the NeuronJ plugin (36) for Fiji/ImageJ

### Western blots

Protein extraction from cells using RIPA buffer and western blotting was done as described in (34). Primary antibodies used in western blotting were FOXA1 (Abcam, Ab23738) and GAPDH (Millipore, MAB374). Immunoreactive bands were visualised using Clarity Western ECL Substrate (Bio-Rad).

### Organoid culture and transfections

PDXs were established by the Melbourne Urology Research Alliance (MURAL) (Monash University Human Research Ethics Committee approval 12287). The established PDXs were grown as subcutaneous grafts in male NSG mice supplemented with testosterone implants according to animal ethics approval (17963), as previously described (33,37). PDXs were routinely authenticated using short tandem repeat profiling (GenePrint 10, Promega) at the Australian Genome Research Facility. Tissue from PDXs 201.1 dura (adenocarcinoma) and 201.2 lung (AR-null) was digested and grown as organoids in growth factor reduced, phenol red-free, ldEV-free Matrigel (Corning). 201.1 organoids were cultured in advanced DMEM/F-12 media (Gibco) containing 0.1 mg/ml Primocin (Invivogen), 1x Glutamax (Gibco), 10 mM HEPES (Gibco), 1 nM DHT (Sigma), 1.25mM N-acetylcysteine (Sigma), 5nM NRG1 Heregulinβ-1 (Peprotech), 500 nM A83-01 (Sigma), 10 mM nicotinamide (Sigma), 0.5 μM SB202190 (Sigma), 2% B27 (Thermo), 20 ng/ml FGF10 (Peprotech), 5 ng/ml FGF7 (Peprotech), 10ng/ml Amphiregulin (Peprotech), 1 μM prostaglandin E2 (Tocris), 10% noggin conditioned media and 10% R-spondin conditioned media. 201.2 Lung organoids were cultured in PrENR -p38i -NAC media (38). 10 μM Y-27632 dihydrochloride (Selleck Chemicals) was added to culture medium during organoid establishment and following passage.

Phase contrast images of organoids were obtained with a Leica DM IL LED microscope with Leica DFC425 C digital camera. For immunohistochemistry, organoids were pelleted in agar, then formalin-fixed and paraffin embedded. Sections were stained using a Leica BOND-MAX-TM autostainer with BondTM epitope retrieval 1 or 2 and the Bond Refine Detection Kit (Leica). Primary antibodies are listed in Supplementary Table S1.

Organoids were transfected with miR LNA inhibitors essentially as described previously (39). Briefly, organoid were collected and 50,000 cells were resuspended in 450µl of organoid culture media and 50μl of transfection mix containing RNAiMAX with 25, 100, and 250 nM miR-194 or NC LNA inhibitor. Cells were centrifuged in a pre-warmed centrifuge at 32°C, 600g for 1h. After centrifuging, cells were incubated in a tissue culture incubator at 37°C for 2-4h and then collected in 1.5ml centrifuge tubes by centrifugation at 300 g for 5 min at room temperature. Cell pellets were resuspended in 50 ul Matrigel and seeded out in 10 μl matrigel discs in 96-well plates. Plates were inverted and incubated at 37°C for 15 minutes to allow Matrigel to solidify, then overlaid with 100mL of organoid culture medium. Organoid forming efficiency was determined at 7 days post-transfection as described previously (33). Organoid viability was assessed at 7 days post-transfection using the CellTiter-Glo® Luminescent Cell Viability Assay kit (Promega), as per the manufacturer’s instructions.

### Statistical Analysis

Statistical analysis for grouped quantitative data were carried out using two-tailed unpaired t-test or ANOVA (GraphPad Prism 7). The relationships between activity scores were determined using Pearson’s correlation coefficient (Graphpad Prism 7).

## RESULTS

### Global identification of transcripts targeted by miR-194 in prostate cancer

Our earlier work demonstrated that miR-194 can promote epithelial-mesenchymal transition (EMT) and metastasis, at least in part by targeting the tumour suppressor SOCS2 (32). However, miRNAs target tens to hundreds of genes, so we hypothesised that elucidating additional miR-194 targets would shed further light on its oncogenic functions in PCa. Thus, we performed Ago-HITS-CLIP on control- and miR-194-transfected 22Rv1 cells to decode miRNA-mRNA interactions. The 22Rv1 model was chosen for this experiment since it exhibits increased metastatic capacity upon transient delivery of miR-194 (32). After immunoprecipitation of Argonaute, co-immunoprecipitating RNA was isolated and evaluated by high-throughput sequencing. Argonaute binding sites (i.e. peaks) that were enriched in cells transfected with miR-194 compared to control transfected cells were identified using MACS2 (17), yielding 7,772 peaks associated with 3,326 genes (Supplementary Table S2). An example peak at the *ZBTB10* gene is shown in Figure 1A. Highlighting the robustness of the data, the vast majority (94%) of peaks were within genes, most commonly in exons, 3’UTRs and introns (Figure 1B). Furthermore, unbiased *de novo* motif analysis revealed that the most strongly enriched sequence within the peaks was a seed recognition site for miR-194 (Supplementary Table S3), which was concentrated within the centres of peaks (Figure 1C).

**Figure 1.**
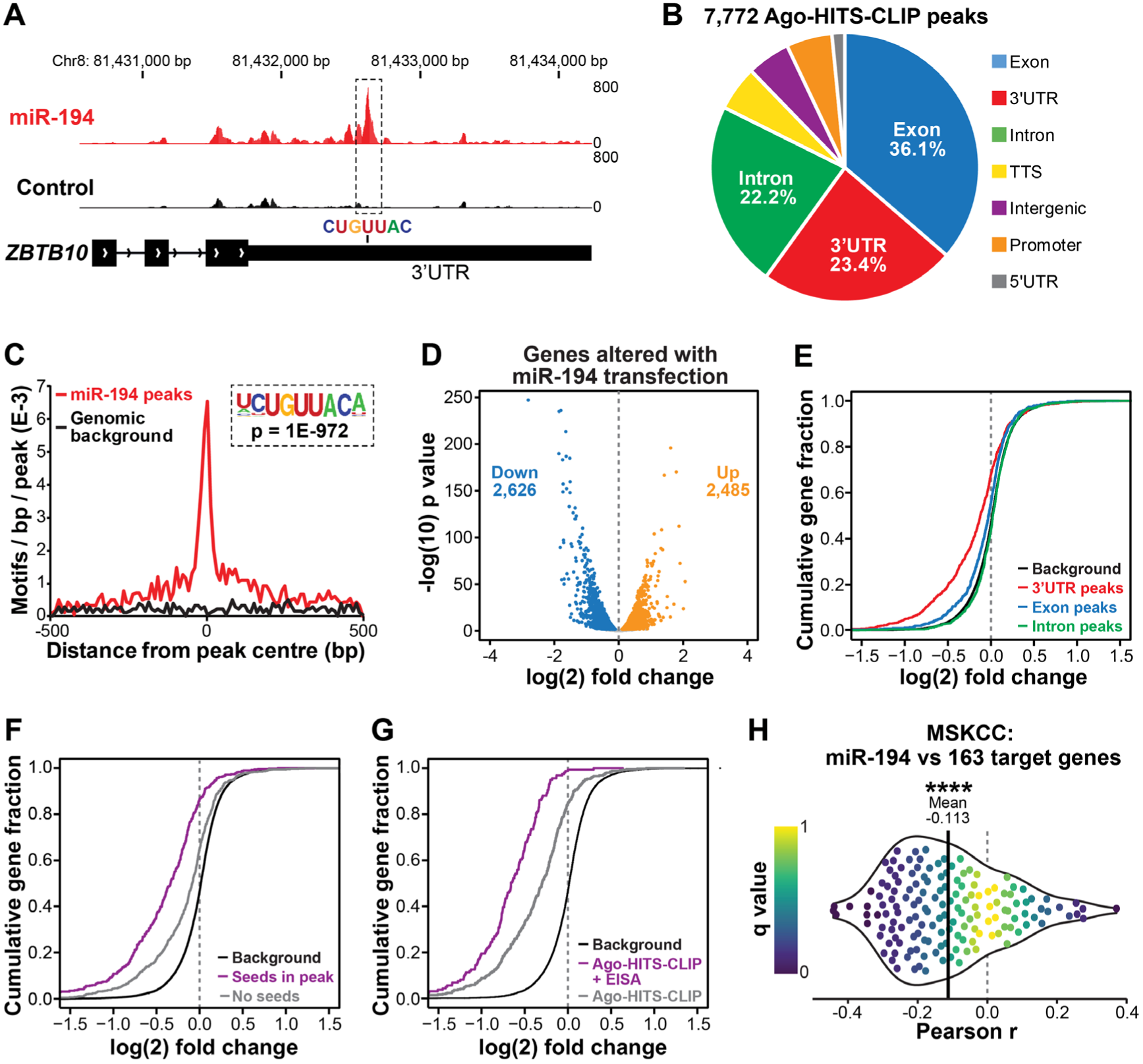
Integrative omics identifies a miR-194 “targetome” in prostate cancer. (A) Example of an Ago-HITS-CLIP peak with a miR-194 seed recognition sequence in the *ZBTB10* gene. Genome tracks depict the average read density of all replicates for each treatment condition (i.e. cells transfected with miR-194 (red) or a scrambled control (black)). (B) Distribution of 7,772 Ago-HITS-CLIP peaks mapped to their genomic regions. (C) Distribution of miR-194 recognition sequences within Ago-HIT-CLIP peaks. Background represents occurrence of the motif on the opposite strand of the peak. (D) Volcano plot showing expression of genes altered by miR-194 transfection in 22RV1 cells. Blue dots indicate significantly downregulated genes and orange dots indicate significantly upregulated genes (FDR ≤ 0.05). (E) Cumulative distribution of log2 fold change for genes containing peaks in the 3’UTR, CDS and Introns compared to a background of all genes with no peaks. (F) Cumulative distribution of log2 fold change for genes with a 3’UTR Ago-HITS-CLIP peak containing miR-194 seed matches or no seed matches in peaks compared to a background of all genes with no peaks. (G) Cumulative distribution of log2 fold change for all genes with a 3’UTR Ago-HITS-CLIP peak containing a miR-194 seed match (Ago-HITS-CLIP) or both a 3’UTR peak with miR-194 seed match and down-regulation at the post-transcriptional level (Ago-HITS-CLIP + EISA). (H) Correlations between miR-194 expression and its “targetome” in 72 primary and metastatic prostate cancers (MSKCC cohort (42)). For each target identified, the Pearson correlation coefficient and its q value was calculated and plotted as -log q (on y-axis) versus correlation coefficient (on x-axis). To indicate the bias towards negative correlations, the mean correlation coefficient is indicated by a vertical black line and the 25th and 75th percentiles are coloured red and blue, respectively. P value was determined using a one-sided t test (****, p < 0.0001).

MiRNAs typically reduce the levels of their target mRNAs (40). Therefore, we conducted RNA-seq of cells transfected with miR-194 mimic or a control. MiR-194 elicited significant changes to the transcriptome, causing down-regulation of 2,626 and up-regulation of 2,485 transcripts (Figure 1D). We identified a strong bias toward down-regulation of mRNAs with miR-194 Ago-HITS-CLIP peaks in 3’UTRs, whereas mRNAs with peaks in coding regions were less strongly biased toward down-regulation and those with peaks in introns were collectively unchanged (Figure 1E; Supplementary Figure S1A). This is consistent with previous studies demonstrating that 3’UTRs are the key sequences through which miRNAs exert their activity (40). Similarly, transcripts with 3’UTR peaks containing miR-194 seed recognition sequences tended to be more robustly down-regulated than those lacking such sequences (Figure 1F; Supplementary Figure S1B).

To further prioritise putative miR-194 target genes, we utilised Exon-Intron Split Analysis, a bioinformatic technique that separates transcriptional and post-transcriptional effects by evaluating whether RNA-seq reads map to exons or introns (41). The quality and number of predicted 3’UTR miR-194 target sites was strongly associated with genes down-regulated at a post-transcriptional but not a transcriptional level (Supplementary Figure S1C), indicating that EISA can differentiate between direct miR-194 targets and downstream changes to the transcriptome. Moreover, transcripts exhibiting post-transcriptional but not transcriptional alterations exhibited stronger down-regulation by miR-194 compared to all transcripts containing Ago-HITS-CLIP peaks (Figure 1G).

With these observations in mind, we applied a filtering strategy whereby transcripts with 3’UTR Ago-HITS-CLIP peaks containing seed recognition sequences and predicted to be down-regulated post-transcriptionally were included in a final 163-gene miR-194 “targetome”. To assess the biological relevance of the “targetome”, its correlation with miR-194 levels was evaluated in 65 primary tumours and 7 metastases (42). We observed a distinct bias toward inverse correlation between miR-194 and its target genes (Figure 1H), supporting the notion that our experimental strategy integrating biochemistry, molecular biology and bioinformatics (i.e. Ago-HITS-CLIP, RNA-seq and EISA) identified *bona fide* targets.

### miR-194 expression and activity is negatively correlated with AR signalling

Gene ontology analysis of the miR-194 targetome revealed enrichment for genes associated with cytoskeletal remodelling, cell adhesion and cell motility (Supplementary Table S4), which likely relates to the ability of miR-194 to enhance prostate cancer cell migration and invasion and elicit an EMT (32). To more specifically evaluate the targetome in clinical prostate cancer, we used single sample gene set enrichment analysis (ssGSEA) of our high-confidence targetome to generate miR-194 activity scores in clinical cohorts, which were then compared to equivalent scores generated from the same cohorts for the “Hallmark” biological gene sets (43). Amongst other robust associations, one striking finding was that miR-194 activity was strongly inversely correlated with AR signalling across all cohorts examined (Figure 2A; Supplementary Table S5). This observation was validated using a more refined set of AR target genes (Figure 2B) recently generated by Sowalsky and colleagues (44).

**Figure 2.**
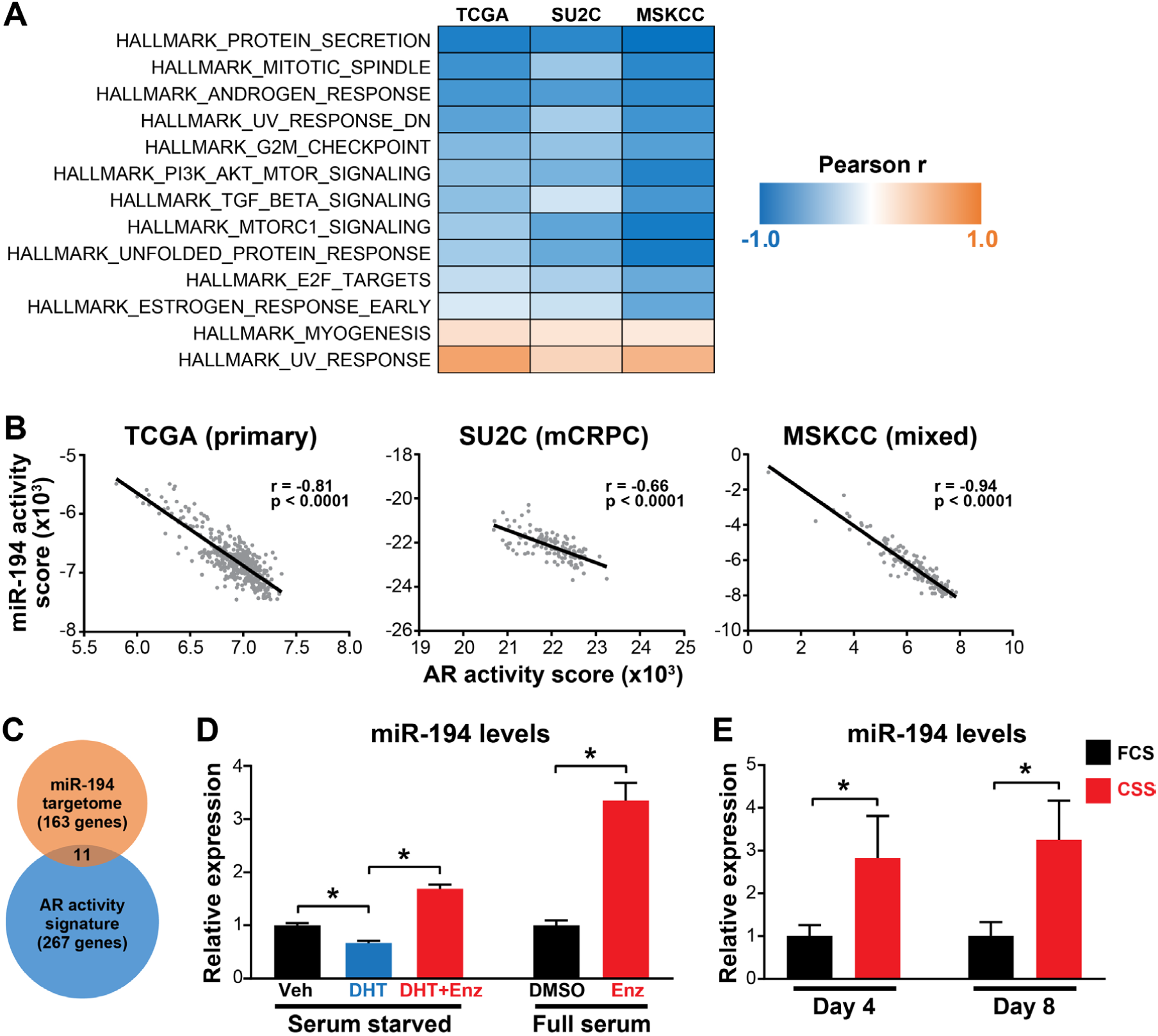
MiR-194 expression is suppressed by AR. (A) Correlation of miR-194 activity score with activity scores of “Hallmark” biological gene sets in the TCGA, SU2C and MSKCC cohorts. P and r values were determined using Pearson’s correlation tests. Only gene sets that were significantly correlated (p < 0.05) in all three cohorts are shown in the heatmap. (B) MiR-194 activity is inversely correlated with AR activity in primary prostate cancer (TCGA cohort, left (65)), metastatic prostate cancer (SU2C cohort, centre (66)) and a cohort comprising both primary and metastatic prostate cancer (MSKCC cohort, right (42)). P and r values were determined using Pearson’s correlation tests. (C) Overlap between the miR-194 targetome and an AR target gene set (44). (D) Relative miR-194 expression in LNCaP cells treated with the androgen DHT and AR antagonist Enzalutamide (Enz). Cells grown in serum starved conditions were treated with vehicle control (Veh) or 10 nM DHT in the presence or absence of 10 μM Enz for 48 hours. Cells grown in full serum were treated with vehicle (DMSO) or 10μM Enz for 48 hours. Expression of miR-194 was normalised to the reference small RNA U6. P values were determined using unpaired two-sided t tests (*, p < 0.05). (E) Relative miR-194 expression in LNCaP cells grown in fetal calf serum (FCS) or charcoal stripped serum (CSS) for 4 or 8 days. Expression of miR-194 was normalised to the reference small RNA U6. P value was determined using an unpaired two-sided t test (*, p < 0.05).

The strength of this negative association led us to examine whether the miR-194 targetome was enriched for AR target genes, but there was only a limited overlap between these gene sets (Figure 2C). Moreover, our Ago-HITS-CLIP and transcriptomic data indicated that miR-194 does not target the *AR* transcript (Supplementary Figure S2). An alternative (and/or additional) explanation for this inverse relationship could be that AR regulates the expression of miR-194. Indeed, levels of miR-194 in the androgen-sensitive LNCaP model were decreased by the potent androgen DHT but increased by the AR antagonist Enzalutamide (Figure 2D). In accordance with these findings, extended culture of cells in androgen-depleted conditions led to upregulation of miR-194 (Figure 2E). Collectively, these data reveal that AR represses expression of miR-194, which (at least partly) explains the negative association between these factors in clinical prostate cancer.

### miR-194 activity and expression is elevated in neuroendocrine prostate cancer

NEPC is associated with loss of canonical AR activity (1). Given the inverse relationship between miR-194 and AR, we therefore hypothesised that its activity would be elevated in clinical NEPC. Indeed, miR-194 activity (estimated by ssGSEA) was significantly higher in NEPC compared to CRPC-Adeno tumours in multiple distinct cohorts (Figure 3A and Supplementary Figure S3). Moreover, miR-194 activity was correlated with established NEPC gene sets (Figure 3B).

**Figure 3.**
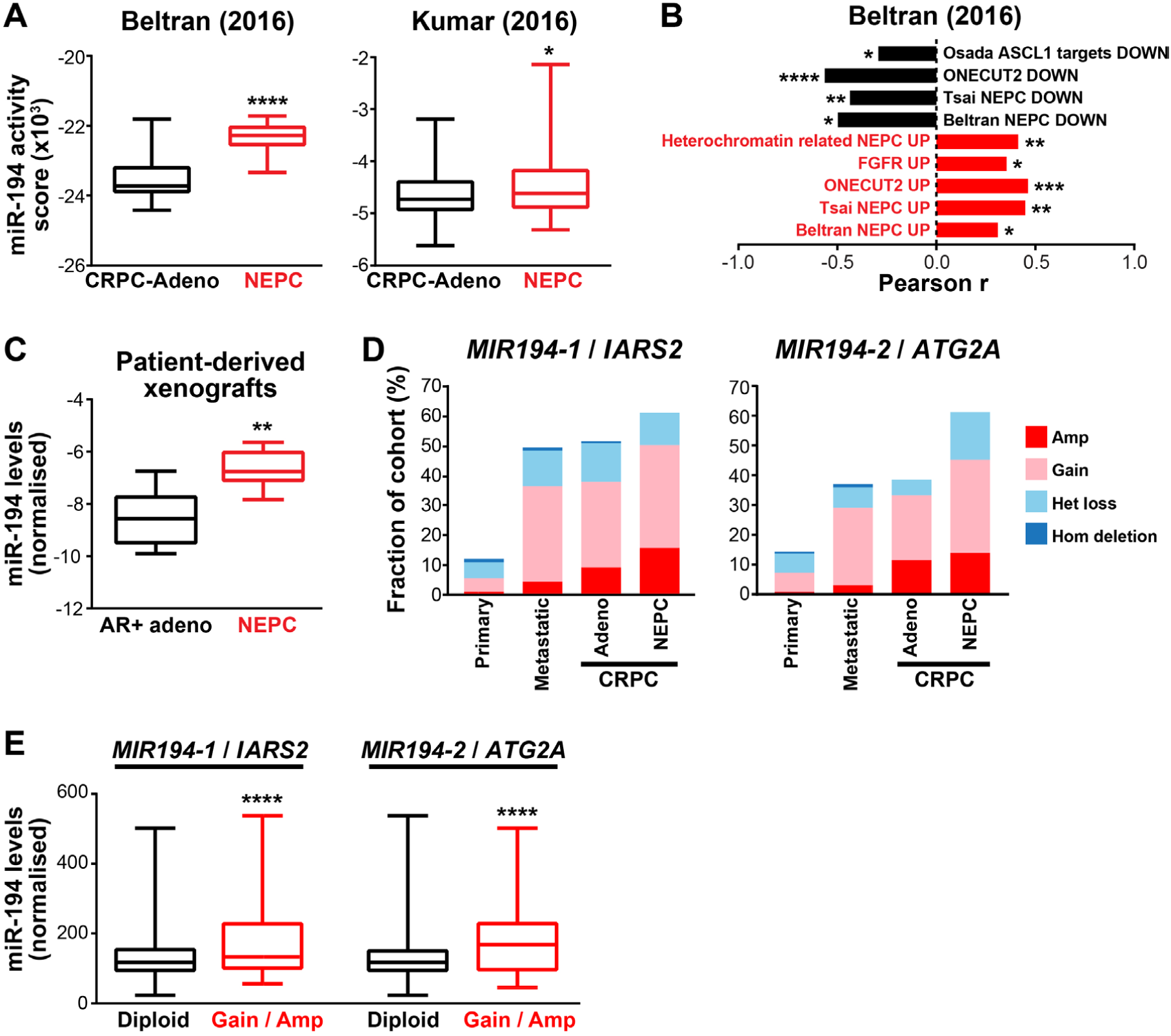
MiR-194 is associated with the AR independent NEPC subtype. (A) MiR-194 activity is higher in neuroendocrine prostate cancer (NEPC) compared to adenocarcinoma CRPC (CRPC-Adeno) in 2 distinct cohorts (6,26). Middle line, mean; lines above and below, ±SD; top and bottom, maximum and minimum. P values were determined using unpaired two-sided t tests (*, P < 0.05; ****, P< 0.0001). (B) Correlation between miR-194 activity and published NEPC associated gene signatures (30,67-71). P and r values were determined using Pearson’s correlation tests (*, p < 0.05; **, p < 0.01; ***, p < 0.001; ****, p < 0.0001). (C) Expression of miR-194 is higher in NEPC PDXs compared to PDXs derived from AR-positive adenocarcinoma tumours. Expression of miR-194 was normalised to two reference small RNAs (U6 and RNU44). Middle line, mean; lines above and below, ±SD; top and bottom, maximum and minimum. P value was determined using an unpaired two-sided t test (**, P < 0.01). (D) *MIR194-1*/*IARS2* and *MIR194-2*/*ATG2A* are more frequently gained/amplified in metastatic compared to primary PCa and in NEPC compared to CRPC-Adeno. Copy number data is combined from multiple clinical cohorts (27,30,72-74). (E) Expression of miR-194 is higher in primary prostate tumours with *MIR194-1*/*IARS2* or *MIR194-2*/*ATG2A* copy number gain or amplification compared to tumours with no change in copy number (diploid). Data is from the TCGA cohort. Middle line, mean; lines above and below, ±SD; top and bottom, maximum and minimum. P values were determined using unpaired two-sided t tests. (****, p<0.0001).

We next examined whether miR-194 itself was over-expressed in NEPC. In the absence of miRNA expression data from clinical samples, we turned to a panel of 13 patient-derived xenografts (PDXs) established through the Melbourne Urological Research Alliance (MURAL), 6 of which have features of NEPC (33). Importantly, miR-194 expression was higher in the NEPC versus AR-positive adenocarcinoma PDXs (Figure 3C), further demonstrating its association with this disease subtype.

Loss of AR expression and/or activity during the transition to NEPC likely explains – at least partly - increased miR-194 expression in this disease state. However, since we have also noted elevated miR-194 expression and activity in metastases and “poor outcome” primary tumours (45), we speculated that other alterations may underlie dysregulation of miR-194 in PCa. MiR-194 is encoded by two separate loci on chromosomes 1 and 11; the *MIR194-1* gene clusters with *MIR215* within intron 12 of the *IARS2* gene on chromosome 1, while the *MIR194-2* gene clusters with *MIR192* approximately 3kb downstream of the *ATG2A* gene on chromosome 11. By interrogating clinical genomic datasets, we found that *MIR194-1*/*IARS2* and *MIR194-2*/*ATG2A* are more frequently gained/amplified in metastatic compared to primary PCa and in NEPC compared to CRPC-Adeno (Figure 3D). Importantly, gain/amplification of these loci were associated with elevated miR-194 levels (Figure 3E). These data suggest that copy number gain can result in increased miR-194 expression in aggressive prostate tumours and NEPC.

### miR-194 promotes the emergence of a neuroendocrine features in prostate cancer

To determine whether miR-194 can directly influence the emergence of a NE-like state, we examined the response of adenocarcinoma PCa cells to transfection with a miR-194 mimic. Exogenous miR-194 led to upregulation of NE marker genes (Figure 4A) and increased neurite length in LNCaP cells (Figure 4B), an effect that was recapitulated in the 22Rv1 cell line model (Figures 4A-B, Supplementary Figure S4).

**Figure 4.**
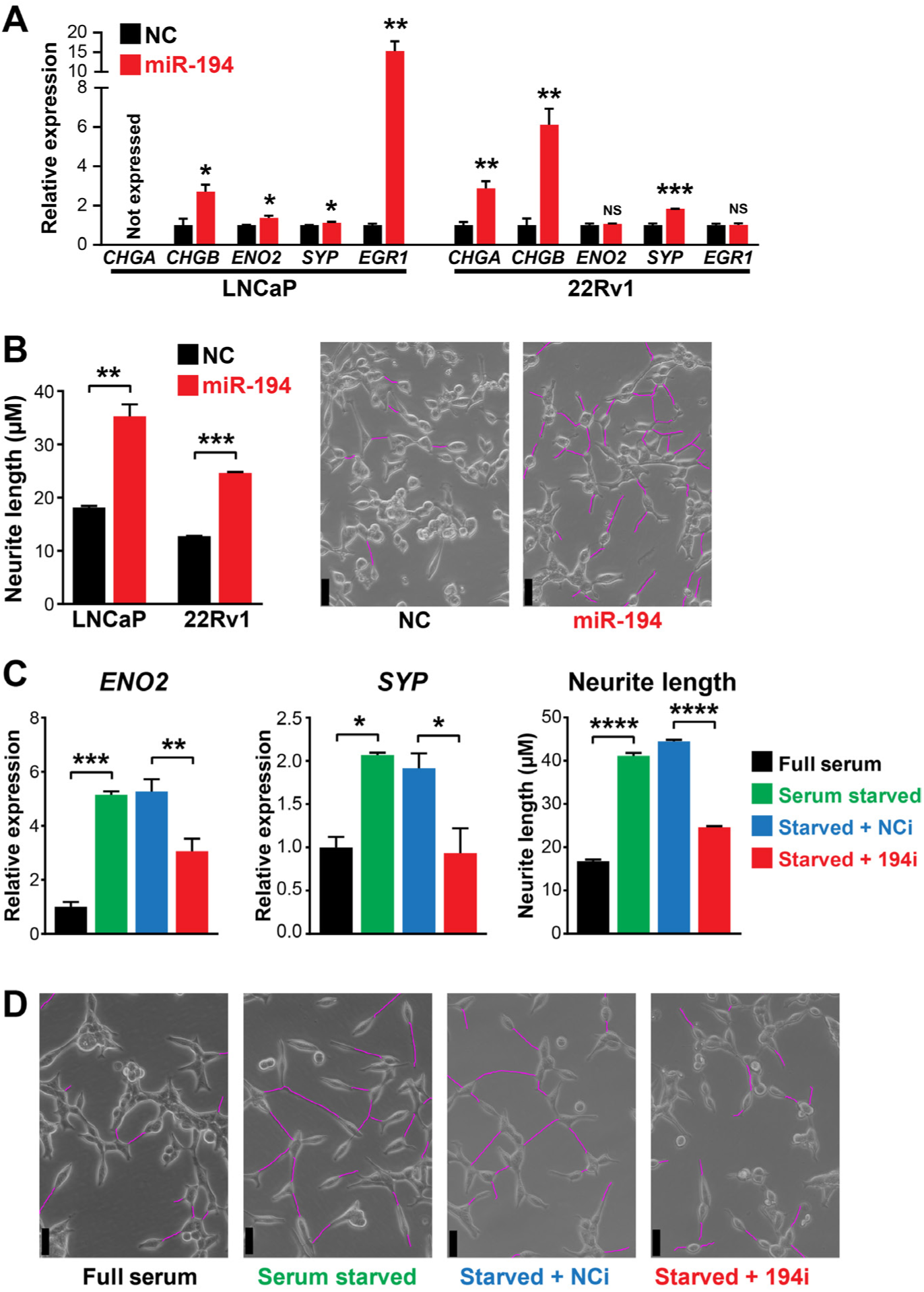
MiR-194 promotes prostate cancer transdifferentiation. (A) Expression of NEPC marker genes is upregulated in response to transfection of a miR-194 mimic in LNCaP and 22RV1 cells. Gene expression was normalised to GAPDH. Expression for the negative control (NC) was set to 1, and error bars are SEM. (B) MiR-194 increases neurite length in LNCaP and 22RV1 cells compared to cells transfected with a negative control miRNA mimic (NC). Expression for NC was set to 1, and error bars are SEM. P values were determined using unpaired two-sided t tests (**, p < 0.01). Representative phase contrast images (on the right) are of LNCaP cells transfected with miR-194 mimic or NC. Neurite outgrowths are traced on images in magenta. Scale bars, 25 μm. (C) A miR-194 inhibitor (194i) blocks neuroendocrine transdifferentiation of LNCaP cells mediated by androgen deprivation, as determined by expression of Neuron specific enolase (*ENO2*), Synaptophysin (*SYP*) and changes in neurite length. Gene expression was normalised to GAPDH. Gene expression or neurite length for cells grown in full serum were set to 1, and error bars are SEM. P values were determined using ANOVA (*, p < 0.05; **, p < 0.01; ***, p < 0.001; ****, p < 0.0001). (D) Representative phase contrast images of LNCaP cells grown in full or stripped serum conditions with or without a miR-194 inhibitor (194i) or negative control inhibitor (NCi). Neurite outgrowths are traced in magenta. Scale bars, 25 μm.

The ability of miR-194 to enhance NE transdifferentiation was further tested using a locked nucleic acid (LNA) inhibitor that specifically inhibits the activity of this oncogenic miRNA. In these experiments, we exploited the fact that the LNCaP model can be transdifferentiated from adenocarcinoma-like to NE-like cells by androgen deprivation (46). As expected, growth of cells in charcoal-stripped serum (CSS) resulted in upregulation of NE markers *ENO2* (encoding neuron-Specific Enolase) and *SYP* (encoding synaptophysin) and increased the length of neurite-like extensions (Figures 4C-D). Importantly, the miR-194 LNA inhibitor effectively blocked this transdifferentiation (Figures 4C-D). Collectively, these data reveal that miR-194 can drive the acquisition of NE features, which corresponds with its increased activity in clinical NEPC.

### FOXA1, an inhibitor of neuroendocrine transdifferentiation, is targeted by miR-194

To understand at a mechanistic level how miR-194 promotes PCa transdifferentiation, we searched the targetome for genes with a known role in PCa progression. Of particular interest was FOXA1, a transcription factor with a critical role in maintaining epithelial lineage in the prostate (47). Consistent with this function, a recent report demonstrated that loss of FOXA1 leads to NE differentiation in prostate cancer (48). Multiple miR-194 Ago-HITS-CLIP peaks were found within the *FOXA1* 3’UTR, one of which contains a perfect 7-mer seed match (Figure 5A). We confirmed that miR-194 decreases the levels of *FOXA1* mRNA and FOXA1 protein in the LNCaP and 22Rv1 models (Figures 5B-C). Importantly, the activity of FOXA1 and miR-194 is inversely correlated in clinical PCa (NEPC and primary PCa; Figure 5D). Collectively, these findings reveal a functional interaction between miR-194 and FOXA1.

**Figure 5.**
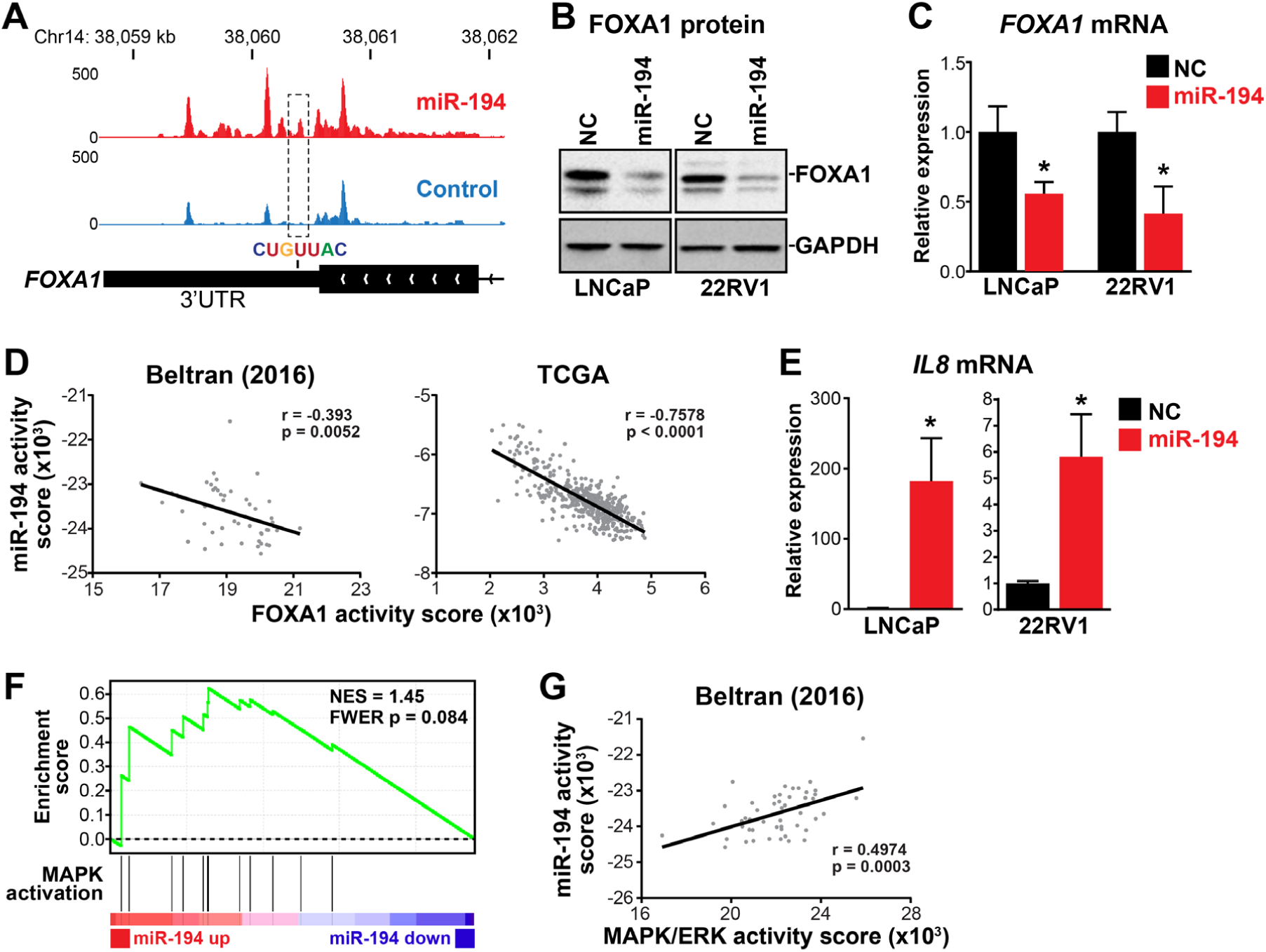
MiR-194 targets *FOXA1* and activates the MAPK/ERK pathway. (A) Ago-HITS-CLIP peaks (including one indicated with a miR-194 seed recognition) sequence in the *FOXA1* gene. Genome tracks depict the average read density of all replicates for each treatment condition (i.e. cells transfected with miR-194 (red) or a scrambled control (black)). (B) Western blot showing FOXA1 protein levels following transfection of a miR-194 mimic or negative control mimic (NC) in 22RV1 and LNCaP cells. (C) Expression of *FOXA1* mRNA, as determined by qRT-PCR, following transfection of miR-194 mimic or NC in 22RV1 and LNCaP cells. Gene expression was normalised to GAPDH. Expression for NC was set to 1, and error bars are SEM. P values were determined using unpaired two-sided t tests (*, p < 0.05). (D) FOXA1 activity is negatively correlated with miR-194 activity in clinical cohorts (6,65). P and r values were determined using Pearson’s correlation tests. (E) Expression of *IL8* is upregulated in response to miR-194 in 22RV1 and LNCaP cells. Gene expression was normalised to GAPDH. Expression for NC was set to 1, and error bars are SEM. P values were determined using unpaired two-sided t tests (*, p < 0.05). (F) MiR-194 causes increased MAPK activity, as determined by GSEA. The MAPK activation signature has been described previously (75). (G) MAPK/ERK activity is positively correlated with miR-194 activity in a clinical cohort comprised of NEPC and CRPC-Adeno samples (6). P and r values were determined using a Pearson’s correlation test.

A recently described mechanism by which FOXA1 suppresses NE differentiation is by directly repressing the *IL8* gene, a chemokine elevated in NEPC, which results in dampening of the MAPK/ERK pathway, a known driver of NEPC (48,49). Supporting the relevance of this mechanism in miR-194’s mode of action, transfection of prostate cancer cells (LNCaP and 22Rv1) with miR-194 caused upregulation of *IL8* (Figure 5E). Moreover, miR-194 also enhanced MAPK/ERK pathway activity (Figure 5F), and miR-194 activity was positively correlated with MAPK/ERK gene signatures in clinical NEPC (Figure 5G and Supplementary Figure S5). Collectively, these data reveal that miR-194 promotes the emergence of NEPC at least in part by targeting FOXA1, which leads to upregulation of IL8 and enhanced MAPK/ERK pathway activity.

### Targeting miR-194 suppresses the growth of prostate cancer with neuroendocrine features

Although miR-194 mediates the acquisition of an NE-like phenotype in prostate cancer, whether it represented a therapeutic target in this disease context is unclear. To investigate this possibility, we measured the growth of PCa cells treated with the miR-194 LNA inhibitor. Interestingly, the growth of all 4 cell line models tested could be suppressed by the inhibitor, but the models with NE features (PC3 and LNCaP-MR42D) were more sensitive than those with a more typical adenocarcinoma phenotype (LNCaP, LNCaP-MR49F) (Figure 6A). The miR-194 inhibitor was cytotoxic as revealed by cell viability assays (Figure 6B) and by counting dead cells (Figure 6C).

**Figure 6.**
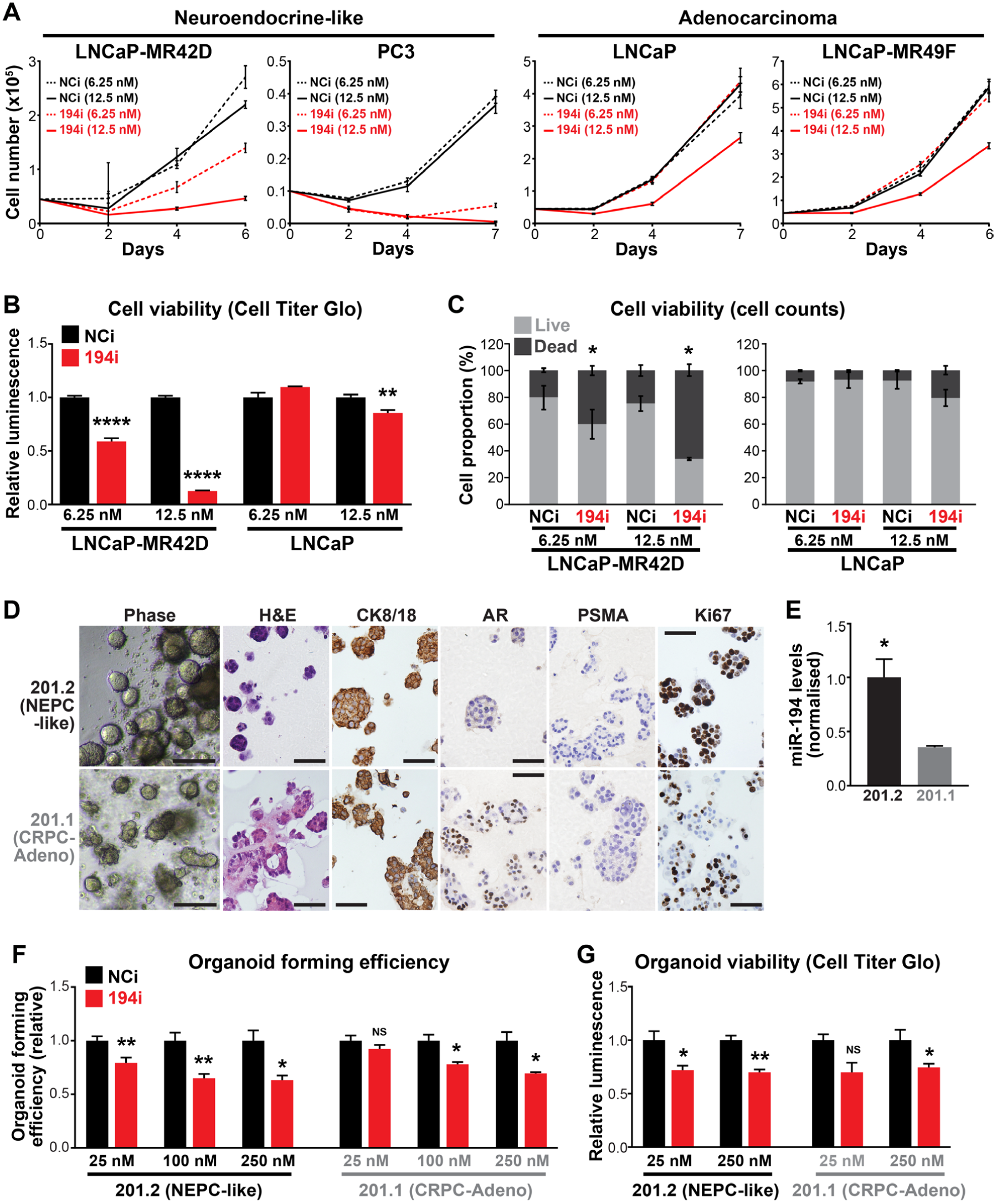
Inhibiting miR-194 blocks the growth of NEPC. (A) Blocking miR-194 activity with an LNA inhibitor (miR-194i) supresses the growth of cell lines with neuroendocrine features (LNCaP-MR42D, PC3) more potently than AR-driven adenocarcinoma cell lines (LNCaP-MR49F, LNCaP), as determined by Trypan blue growth assays. Error bars are SEM. Negative control inhibitor, NCi. (B) Blocking miR-194 activity with an LNA inhibitor supresses cell viability more potently in neuroendocrine-like LNCaP-MR42D cells compared to LNCaP adenocarcinoma cells, as determined by Cell Titer Glo cell viability assay. P values were determined using unpaired two-sided t tests (**, p < 0.01; ****, p < 0.0001). (C) Proportion of live and dead cells in LNCaP-MR42D and LNCaP cells transfected with 194i or NCi. P values were determined using unpaired two-sided t tests (*, p < 0.05). (D) Representative phase contrast, haematoxylin and eosin (H&E) and immunohistochemistry (IHC) of 201.1 and 201.2 organoid models growing as colonies in Matrigel. Scale bars: phase images = 100 μm; H&E and IHC = 50 μm. (E) Levels of miR-194 in 201.1 and 201.2 organoids. Expression of miR-194 was normalised to 2 reference small RNAs (U6 and RNU44). P value was determined using an unpaired two-sided t test (*, p < 0.05). (F-G) Blocking miR-194 activity with 194i inhibits organoid forming efficiency (F) and organoid viability (G) of the 201.1 and 201.2 models. P values were determined using unpaired two-sided t test. (*, p < 0.05; **, p < 0.01).

To examine the potential of targeting miR-194 in a more clinically-relevant setting, we turned to patient-derived CRPC organoids recently described by our team (33). Models 201.1 and 201.2 were derived from dura and lung metastases, respectively, from a patient who died after receiving second-generation AR-targeted therapies (Enzalutamide, Abiraterone) and chemotherapies (Docetaxel, Cabazitaxel) (33). 201.1 is a model of PSA-positive adenocarcinoma that expresses a mutated form of the AR (C687Y and T878A) that mediates resistance to Enzalutamide. In contrast, 201.2 has no AR or PSA expression but exhibits high expression of a neuroendocrine gene signature, focal staining of CD56, and concurrent genomic loss of *TP53, PTEN* and *RB1* (33). Representative IHC images of the expression of various markers in each model are shown in Figure 6D. As expected, miR-194 levels were higher in 201.2 compared to 201.1 (Figure 6E). The effect of the miR-194 LNA inhibitor on the growth of these 2 patient-derived models was evaluated by measuring organoid forming efficiency (OFE) and cell viability. Both models exhibited reduced OFE and cell viability in response to transfection with the inhibitor (Figures 6F-G). However, similarly to the cell lines, the AR-null, NEPC-like 201.2 model was more sensitive to miR-194 inhibition than the adenocarcinoma-like 201.1 model (Figures 6F-G). Collectively, these findings - in both traditional cell lines and contemporary patient-derived models - provide evidence that targeting miR-194 has potential as a novel therapy for prostate cancer with NE features.

## DISCUSSION

Epigenetic and transcriptional alterations are known to mediate prostate cancer cell plasticity during adenocarcinoma-neuroendocrine transdifferentiation. Most drivers of these alterations identified to date are transcription factors and chromatin modifiers, such as SOX2 (50,51), EZH2 (50), REST (52), BRN2 (8) and FOXA2 (53). By identifying miR-194 as a mediator of this transdifferentiation, our work reveals that post-transcriptional gene regulation is another mechanism by which transcriptional networks are altered during progression to NEPC.

Our study suggests that miR-194 is elevated in NEPC via 2 key mechanisms. First, by evaluating PCa cells treated with androgens and anti-androgens, we found that miR-194 is negatively regulated by the AR signalling axis. Interrogation of published cistromic data revealed no evidence for association of AR with regulatory elements proximal to *MIR194* genes (data not shown); thus, we do not believe that the inverse relationship between AR and miR-194 represents a direct mode of transcriptional repression. Rather, we hypothesise that AR indirectly inhibits miR-194 expression through a mechanism that is yet to be elucidated. One possibility is that the transcription factor GATA2 serves as an intermediary: expression of GATA2 is known to be down-regulated by AR (54), and we previously demonstrated that GATA2 enhances the levels of miR-194 (32). Future studies should investigate the role of this putative pathway in NEPC, particularly since GATA2 has been identified as a mediator of PCa metastasis and drug resistance previously (55). Second, we found that gain or amplification of genomic regions encompassing the *MIR194* genes is another mechanism that can result in elevated expression of miR-194 in aggressive forms of prostate cancer, including NEPC. MiR-194 is unusual in that it is encoded by 2 genes (*MIR194-1* and *MIR194-2*), and the observation that both are frequently gained further supports the relevance of this miRNA in disease progression.

Using an integrative approach that exploited cutting-edge biochemical (Ago-HITS-CLIP), molecular (RNA-seq) and bioinformatics (EISA) techniques, we identified ∼160 genes that miR-194 putatively targets in PCa. Of note, gene signatures enriched in this targetome include those involved in cell movement, cytoskeletal organisation (including axon guidance) and focal adhesion. We propose that dysregulation of these networks by elevated miR-194 during PCa progression EMT (32) and transdifferentiation from an adenocarcinoma-like cell to an NE-like cell (this study). While this hypothesis remains to be proven, we note the EMT and emergence of NE features are manifestations of cell plasticity that share many fundamental characteristics; indeed, it appears as if the re-activation of a developmental EMT program is a crucial strategy by which PCa cells evolve towards a NE lineage (1,56).

In addition to a miR-194 targetome enriched for cell movement, structure and attachment, we identified *FOXA1* as a key target gene via which miR-194 influences the emergence of NEPC. Supporting our findings, *FOXA1* has been previously identified as a target of miR-194 in non-small cell lung cancer (NSCLC); interestingly, in this context it appeared to act as a tumour suppressor, with upregulation of miR-194 suppressing tumour proliferation, invasion and metastasis (57). The divergent outcomes of targeting FOXA1 by miR-194 in PCa and NSCLC reflects a common phenomenon in miRNA biology whereby context-dependent roles are mediated by the relative expression of key miRNA target genes in a particular cell or tissue environment. Our data suggest that targeting of FOXA1 by miR-194 in PCa leads to de-repression of IL8 and subsequent upregulation of the MAPK/ERK pathway (48). Both IL8 and the MAPK/ERK pathway are known drivers of NEPC (49,58,59); our work defines a new mechanism by which these factors are elevated in this disease context.

The relevance of the miR-194:*FOXA1* pathway in PCa likely goes beyond its consequent impact on IL8 and MAPK/ERK, since FOXA1 is a pioneer factor for AR and a major regulator of its transcriptional outputs (60). Like FOXA1, AR is also vital for maintenance of the epithelial phenotype; therefore, the consequent disruption of AR signalling by down-regulation of FOXA1 could be another mechanism by which miR-194 enhances lineage plasticity in PCa. Combined with our finding that miR-194 is repressed by AR signalling and the identification of up to 11 AR downstream genes as miR-194 targets (Figure 2), our study reveals a complex and intimate interplay between miR-194 and this key pathway in PCa. In addition to being the likely explanation for the extremely strong negative correlation between miR-194 and AR in clinical PCa, this interplay may also influence response to AR-targeted therapies.

Given the increasing frequency of treatment-emergent NEPC tumours and their aggressiveness, the development of therapies that selectively target this CRPC subtype is critically important. Indeed, strategies to target AURKA (which promotes the activity of MYCN, a known driver of NEPC), EZH2 (which enhances adenocarcinoma-NEPC transdifferentiation) and the Wnt and NOTCH pathways (both of which promote stem cell maintenance in NE-like tumours) are being evaluated in clinical trials (1). Our study identifies miR-194 as a novel therapeutic target in this disease setting. Although a recent study found that miR-652 can promote the acquisition of NE features in PCa cells (61), to our knowledge ours is the only study to date demonstrating that targeting a miRNA can inhibit NE transdifferentiation and block the growth of NEPC. Moreover, the sensitivity of patient-derived CRPC organoids and PCa cell lines to nanomolar doses of a miR-194 inhibitor highlights the potential of such a therapeutic strategy. While miRNA-based therapies have proven difficult to translate to the clinic (62), at least 2 antagomiRs are currently being evaluated in trials: a miR-122 antagomiR (“Miravirsen”) showed activity in a phase IIa trial of hepatitis C (in which no adverse side effects were reported), while a miR-155 antagomiR is in phase I trials for lymphoma (63). The attraction of targeting miRNAs in cancer comes from the potential to concurrently modulate multiple pathways involved in tumour growth and progression. In the case of miR-194, an inhibitor could stabilise multiple plasticity suppressing factors (e.g. FOXA1) and tumour suppressors (e.g. SOCS2 (32)), leading to inhibition of multiple plasticity- and metastasis-promoting pathways (e.g. MAPK/ERK, IL8 and STAT3). We aim to undertake further pre-clinical evaluation of a miR-194-targeted therapy to treat NEPC and/or re-sensitise NEPC tumours to AR-targeted therapies.

In addition to its potential as a therapeutic target, it is worth noting that miR-194 was first linked to PCa as a serum marker of poor prognosis in a patients with localised disease (45). In this earlier disease context, high levels of serum miR-194 likely demarcates tumours with increased plasticity and hence a propensity to metastasize. However, whether miR-194 is a marker of advanced PCa and CPRC is unknown. Given the strong inverse correlation between miR-194 and AR activity, it is tempting to speculate that circulating miR-194 could be used to identify CRPC patients with AR-independent tumours (e.g. NEPC) and therefore guide therapy, but this concept remains to be tested in patient cohorts.

In summary, our study demonstrates that miR-194 can promote adenocarcinoma-NE transdifferentiation and the growth of NEPC by targeting a network of genes including the lineage-defining transcription factor FoxA1. These findings deliver new molecular insights into lineage plasticity in PCa, and provide impetus to further investigate the potential of targeting miR-194 as a novel therapy for NEPC.

## ACCESSION NUMBERS

Ago-HITS-CLIP and RNA-seq data have been deposited with the Gene Expression Omnibus bank under accession number GSE137072.

## ACKNOWLEDGEMENTS

The authors thank: the Australian Cancer Research Foundation (ACRF) Cancer Genomics Facility for assistance with Ago-HITS-CLIP and RNA-seq; Nicholas Choo (Monash University), Birunthi Niranjan (Monash University), Roxanne Toivanen (Monash University) and Susan Woods (University of Adelaide) for expert technical assistance with organoid culture; Jindan Yu (Northwestern University) for providing a FOXA1 activity gene set; Peter Nelson and Ilsa Coleman (Fred Hutchinson Cancer Research Center) for providing transcriptomic data (64); and Emily Hackett-Jones (University of South Australia) for assistance with analysis of the Ago-HITS-CLIP data. We acknowledge the team that generated transcriptomic data from CRPC-Adeno and NEPC tumours (6), which we obtained from dbGaP (accession number phs000909). The results published here are in part based on data generated by The Cancer Genome Atlas, established by the National Cancer Institute and the National Human Genome Research Institute, and we are grateful to the specimen donors and relevant research groups associated with this project. Finally, we thank the patients and clinicians who have generously supported the MURAL research platform.

