## Supplementary Figures for "MicroRNA-194 promotes lineage plasticity in advanced prostate cancer"

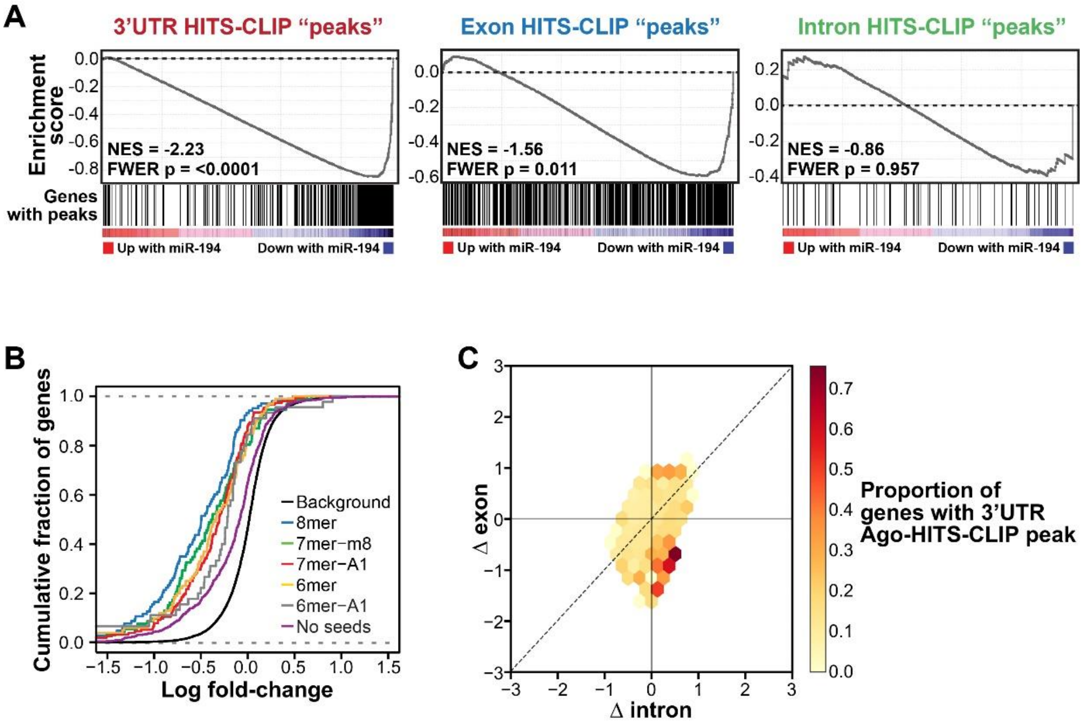


**Supplementary Figure S1. Defining a miR-194 targetome.** (A) Ago-HITS-CLIP peaks in the 3'UTR of genes are associated with their down-regulation, as determined by GSEA. Genes with peaks in exons were biased towards down-regulation, whereas genes with peaks in introns were not collectively down-regulated. (B) Cumulative distribution of log2 fold change for 3'UTR Ago-HITS-CLIP peaks containing different miR-194 seed recognition sequences or no seeds, compared to a background of all genes with no peaks. (C) Post-transcriptionally downregulated genes will have a negative delta exon (Δ exon) and a positive/no delta intron (Δ intron) or a Δ exon-Δ intron < 0. Post-transcriptionally downregulated genes are mainly concentrated in the bottom right quadrant and close to the y-axis in the bottom left quadrant. Genes that have a negative Δ exon but also a negative Δ intron (such that Δ exon-Δ intron = ≥ 0) are likely to be downregulated at the transcriptional level. The colour of the hexamers indicates percentage of genes with Ago-HITS-CLIP peaks in the 3'UTR.

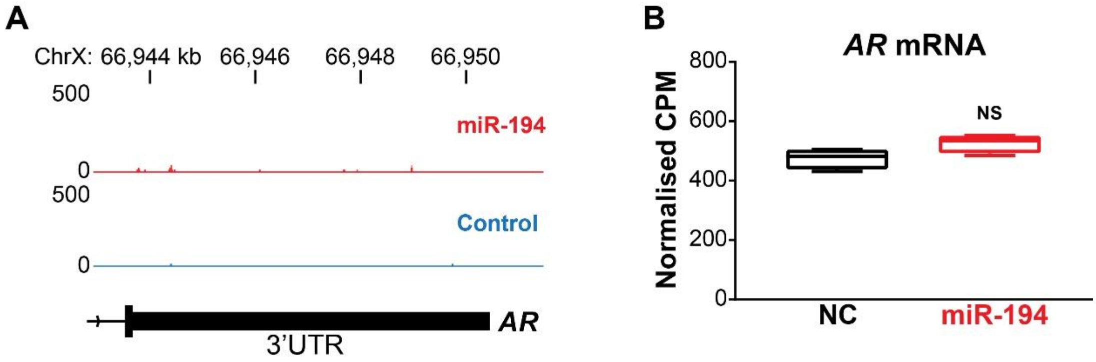


**Supplementary Figure S2. No evidence for direct targeting of *AR* by miR-194.** (A) Genome tracks from the Ago-HITS-CLIP experiment depicting the average read density of all replicates for each treatment condition (i.e. cells transfected with miR-194 (red) or a scrambled control (black)) at the *AR* 3'UTR. No Ago-HITS-CLIP peaks are evident. (B) *AR* mRNA expression is not altered with miR-194 transfection, as determined by RNA-seq.

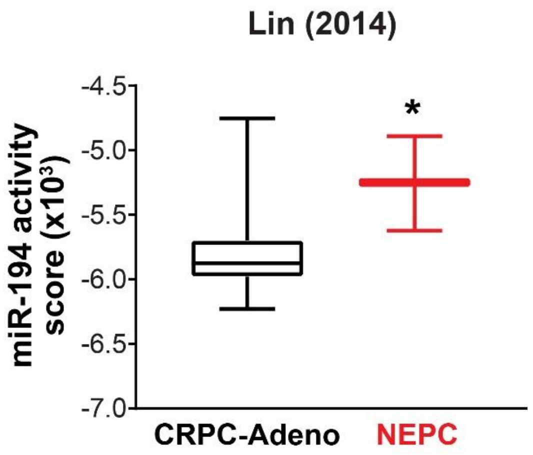


**Supplementary Figure S3.** MiR-194 activity is higher in neuroendocrine prostate cancer (NEPC) patient-derived xenografts (PDXs) compared to adenocarcinoma CRPC (CRPC-Adeno) PDXs (GSE41192) (Lin et al 2014).

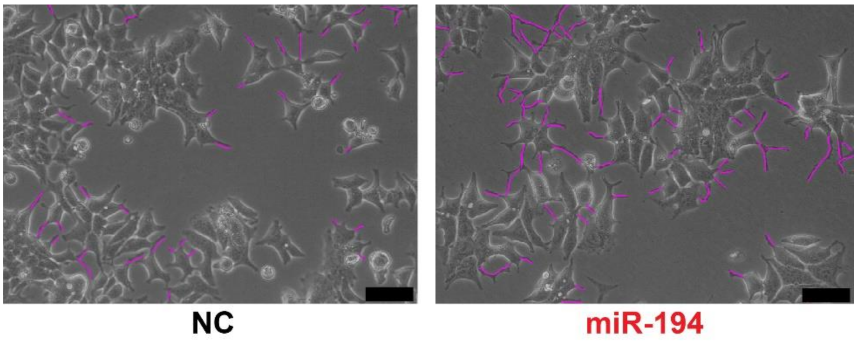


**Supplementary Figure S4.** Representative phase contrast images of 22Rv1 cells transfected with a miR-194 mimic or a negative control (NC) mimic. Neurite outgrowths are traced in magenta. Scale bars, 25 µm.

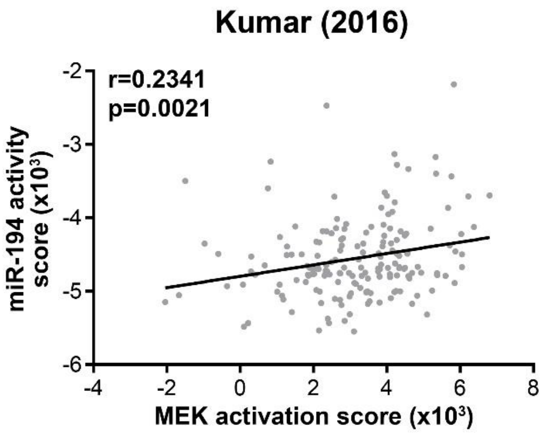


**Supplementary Figure S5.** MiR-194 activity is positively correlated with MAPK pathway activation in a clinical cohort prostate cancer (Kumar et al 2016).
